# Robust inducible gene expression in intracellular *Listeria monocytogenes in vivo*

**DOI:** 10.1101/2024.05.28.596178

**Authors:** Huong Giang Pham, Kiet N. Tran, Larissa Gomelsky, Tathagato Roy, Jason P. Gigley, Mark Gomelsky

## Abstract

Attenuated strains of the intracellular pathogen *Listeria monocytogenes* can deliver genetically encoded payloads inside tumor cells. *L. monocytogenes* preferentially accumulates and propagates inside immune-suppressed tumor microenvironments. To maximize the payload impact in tumors and minimize damage to healthy tissues, it is desirable to induce payload synthesis when bacteria are eliminated from the healthy tissues but are grown to high numbers intratumorally. Here, we have engineered a tightly controlled gene expression system for intracellular *L. monocytogenes* inducible with a cumin derivative, cumate. Upon cumate addition, expression of a reporter gene is increased in *L. monocytogenes* growing *in vitro* by 80-fold, and in intracellular *L. monocytogenes* in murine tumors by 10-fold. This study demonstrates the feasibility of activating gene expression in intracellular bacteria in live animals using an edible inducer. The system is expected to enhance the efficacy and safety of the attenuated *L. monocytogenes* strains as antitumor payload delivery bacterial drones.

## Introduction

Bacteria-based treatments have emerged as a promising approach in cancer therapy^1^. Bacteria can colonize and thrive in the immune suppressive tumor microenvironments (TMEs), therefore *Clostridium*^2^, *Listeria*^3,4^, *Escherichia*^5,6^, *Salmonella*^7,8^, and others have been engineered as vehicles for tumor-targeted delivery of tumor-associated antigens, foreign antigens, or antitumor payloads. *Listeria monocytogenes* (*Lm*)-based cancer therapy has thus far been most extensively tested in clinical trials^9^. *Lm* is a facultatively intracellular pathogen that stands out due to its ability to strongly trigger both innate and adaptive immune responses in TMEs^9,10^. When injected systemically, *Lm* infects myeloid-derived suppressor cells (MDSCs)^11^, which are actively recruited by tumors. *Lm* hitchhikes inside the MDSCs to the primary tumors and metastases. There, *Lm* spreads from MDSCs to tumor cells^12^. Attenuated, low-virulence *Lm* strains, such as XFL-7, have been used in several clinical trials showing moderate efficacy but acceptable safety^9^. XFL-7 lacks the master transcriptional activator of virulence genes, PrfA, required for *Lm* survival inside mammalian cells^3,13–15^. This deficiency is partially complemented by the plasmid-encoded *prfA* and truncated version of listeriolysin O (*trLLO*)^3,14,15^.

Even attenuated strains that colonize healthy tissues following systemic *Lm* delivery may cause significant damage if they constitutively produce cell-toxic agents^12^. Further, constitutive expression of toxins inhibits bacterial proliferation inside TMEs. Both the safety and efficacy of *Lm*-based cancer therapy can be enhanced if payload synthesis is induced when most *Lm*-infected cells in the healthy tissues are already eliminated and the number of *Lm*-infected cells inside the immune suppressive TMEs is high. However, previously constructed gene regulatory systems in *Lm* are not well-suited for *in vivo* applications due to low inducer cell permeability, toxicity and/or system leakiness^16–20^.

In this study, we developed a cumate-inducible system for robust gene activation in intracellular *Lm* in tumors. This system utilizes the membrane-permeable, low toxicity compound, cumate, 4-isopropylbenzoate. Cumate is derived from cumin, a culinary spice, which makes it a safe candidate for applications in humans. While cumate-inducible systems have previously been demonstrated to regulate gene expression in prokaryotic and eukaryotic microbes as well as in mammalian cells^21–24^, it has remained unclear if cumate can induce expression in intracellular bacteria. In this study, we engineered a tightly cumate-dependent system for inducing genes in intracellular *Lm* localized inside murine tumors.

## Materials and Methods

### Bacteria Strains and Growth Conditions

*E. coli* strains NEB Stable and NEB 10β (NEB) were cultured in Luria–Bertani (LB) media at 37 °C. *Lm* XFL-7 (a gift from Dr. Gravekamp) was cultured in Brain Heart Infusion media (BHI) at 37 °C. Chloramphenicol was added at 25 µg/mL for *E. coli* and 10 µg/mL for *Lm*; kanamycin was added at 50 µg/mL for *E. coli* and *Lm*.

### Plasmid Engineering

Cloning was performed using restriction enzymes and HiFi DNA assembly (NEB). Primers (Eurofins Genomics) and synthetic DNA fragments (Twist Bioscience) used in this study are listed in Table S1. The cumate-inducible system cassette was amplified from plasmid pCT5-bac2.0 (Addgene, #119872; gifted by Dr. Schmidt-Dannert) and cloned into the *E. coli*-*Lm* shuttle-vector pGG-34^13–15^ lacking trLLO, creating pLicu1.0. The PxylR promoter in pLicu1.0 was replaced with P16, P18, and PT17 promoters, resulting in plasmids pLicu1.1, pLicu1.2, pLicu1.3, respectively. For the *in vivo* cumate-inducible system, the PT17-CymR cassette from pLicu1.3 was cloned into the listerial integrative pIMK2 vector^25^. The P_veg_-CuO-sfGFP cassette with a strong terminator (Tz, Table S1) was amplified and cloned into the pGG-34 vector containing trLLO, resulting in pLicu2.0L, where sfGFP is superfolder-GFP^26^. RBS strength was optimized using an RBS Calculator^27^ to create pLicu2.0M (23K) and pLicu2.0H (96K). All plasmids were transformed into *E. coli* NEB Stable or electroporated into *Lm* XFL-7 as described by Monk *et al*.^25^.

### *In vitro* Induction of GFP Expression in *Lm*

*Lm* XFL-7 strain harboring different cumate-inducible system constructs was grown in 5 mL of BHI containing appropriate antibiotics. Cumate (2–50 µg/mL, Sigma #268402) was added at log phase (A_600_, 0.4–0.6) to induce sfGFP expression. 1-mL culture aliquots at 2, 4, 8, and 24 h post-induction was pelleted and washed twice with PBS before measuring A_600_ and GFP fluorescence using the Glomax Multi JR Detection System (Promega).

### Flow Cytometry

*Lm* cultures grown in BHI broth with varying cumate concentrations were pelleted, washed with PBS, and measured using Guava EasyCyte 12HT flow cytometer (Millipore). Data were analyzed and plotted using FlowJo™ v10.8 Software (BD Life Sciences).

### *In cellulo* Induction in *Lm*

The murine breast cancer cells, 4T1 (ATCC, #CRL-2539), were cultured in RPMI-1640 medium (Corning, #10-040-CV,) with 10 % fetal bovine serum (FBS, #SH3007102, Fisher Scientific, MA), 0.5 µg/mL Amphotericin B (Fisher Scientific, #15-290-018,), and 100 U/mL penicillin-streptomycin (Fisher Scientific, #15-140-122,) with 5 % CO_2_ at 37 ºC. Cells were seeded into 96-well black plates (Sigma, #CLS3603) at 2×10^4^ cells per well. *Lm* cultures at log phase were used to infect 4T1 cells at a multiplicity of induction (MOI) of 10 as previously described^28^. *Lm*-infected 4T1 cells were grown in RPMI with 10% FBS and 25 µg/mL gentamicin for 15 h, then treated with 0–100 µg/mL cumate. sfGFP fluorescence was measured at 2, 4, 8, 24, and 48 h post-induction using an *in vivo* imaging system (IVIS) (Kino, Spectral Instruments Imaging); 10-sec exposure, 10 % excitation power.

For microscopy, 4T1 cells were seeded on coverslips in a 24-well plate at 1×10^5^ cells per well. *Lm* infection and induction were performed as described above. At 24 h post-induction, the coverslips were washed with PBS, fixed with 4% paraformaldehyde for 10 min, and washed twice with PBS. The coverslips were mounted onto a glass slide with DAPI-containing medium (Abcam, Ab104139) and imaged using a ZEISS LSM 980 microscope.

### Induction of Gene Expression in Intracellular *Lm in vivo*

Animal experiments were performed in accordance with NIH guidelines and approved by the University of Wyoming Institutional Animal Care and Use Committee (protocol #2023-0068). Tumors were implanted by injecting 1×10^5^ 4T1 cells into the mammary fat pad of 6-week-old female BALB/c mice (Jackson Labs). *Lm* XFL7 (2×10^7^ CFU/mouse) harboring the cumate-inducible system was injected intratumorally when tumors reached 50–100 mm^3^. The sfGFP expression was induced on day 3 post-injection by intraperitoneal (I.P.) injection of 3.0 mg/mouse of water-soluble cumate (System Biosciences, #QM150A-1). Mice were imaged with IVIS or sacrificed at indicated times for *ex vivo* imaging. Tumors were isolated and fixed with 4% paraformaldehyde at 4 °C overnight, sectioned into 60 µm slices using a vibratome (Technical Products International). Tumor tissues were mounted on glass slides with DAPI-containing medium for visualization with a ZEISS LSM 980 microscope.

### Bioinformatics and Statistics

Ribosome binding site strengths were predicted using the RBS calculator^27^ (https://salislab.net/software/predict_rbs_calculator) for *Lm*. Relative induction of *Lm in vivo* was measured in microscopy images using ImageJ with the JacoP plugin^29^. Backgrounds were normalized before calculating the overlap between red and green channels as a percentage of induction, based on Manders’ coefficient as previously described^29^. Statistical analyses were performed using Prism 5 (GraphPad, CA).

## Results and Discussion

The cumate-inducible system relies on the CymR repressor that binds to the CuO operator sequence located upstream of the gene of interest (GOI). Upon cumate binding, CymR affinity to DNA is decreased thus resulting in GOI de-repression (Fig. 1A). We used the module developed for *Bacillus*, pCT5-bac2.0, as the basis for the cumate-inducible system in *Lm*. Howevver, our attempts to clone this module in the listerial therapeutic vector, pGG-34, proved futile due to plasmid instability (Fig. S1A). Most plasmids isolated from in *E. coli* had deletions, while rare plasmids of the expected size contained mutations in either the *cymR* or the *trLLO* genes, suggesting that a combination of CymR and trLLO is deleterious to *E. coli*. To avoid such combination, we cloned the cumate-inducible system in the pGG-34 derivative lacking the *trLLO* gene, designated pLicu1.0 (Fig. 1B). This plasmid proved to be stable in *E. coli* and in *Lm* XFL-7 (Fig. S1B, C), and resulted in a cumate-dependent expression of the GOI, sfGFP. However, sfGFP expression was leaky (Fig. 1C). We hypothesized that CymR levels were insufficient to repress sfGFP transcription in the absence of cumate, and replaced the PxylR promoter driving *cymR* expression in pLicu1.0 with a series of constitutive promoters P16, P18^30^, PT17^20^, resulting in pLicu1.1, 1.2, and 1.3, respectively (Fig. 1B). Among these constructs, pLicu1.3 showed the tightest repression and the broadest dynamic range, i.e., 50-fold increase in sfGFP expression after 24 h at 20 µg/mL cumate (Figure 1C). Flow cytometry results revealed that only 1.8% of uninduced *Lm* cells expressed some sfGFP, whereas the fraction of sfGFP-expressing cells rapidly increased to the 59-64% range at varying levels of cumate at 24 h post-induction. Importantly, cumate showed no detrimental effect on *Lm* growth, even at very high (500 µg/mL) concentrations (Fig. S2). The cumate-inducible system in pLicu1.3 performed similarly to TetR-dependent system recently developed for *Lm*^20^.

**Fig. 1.**
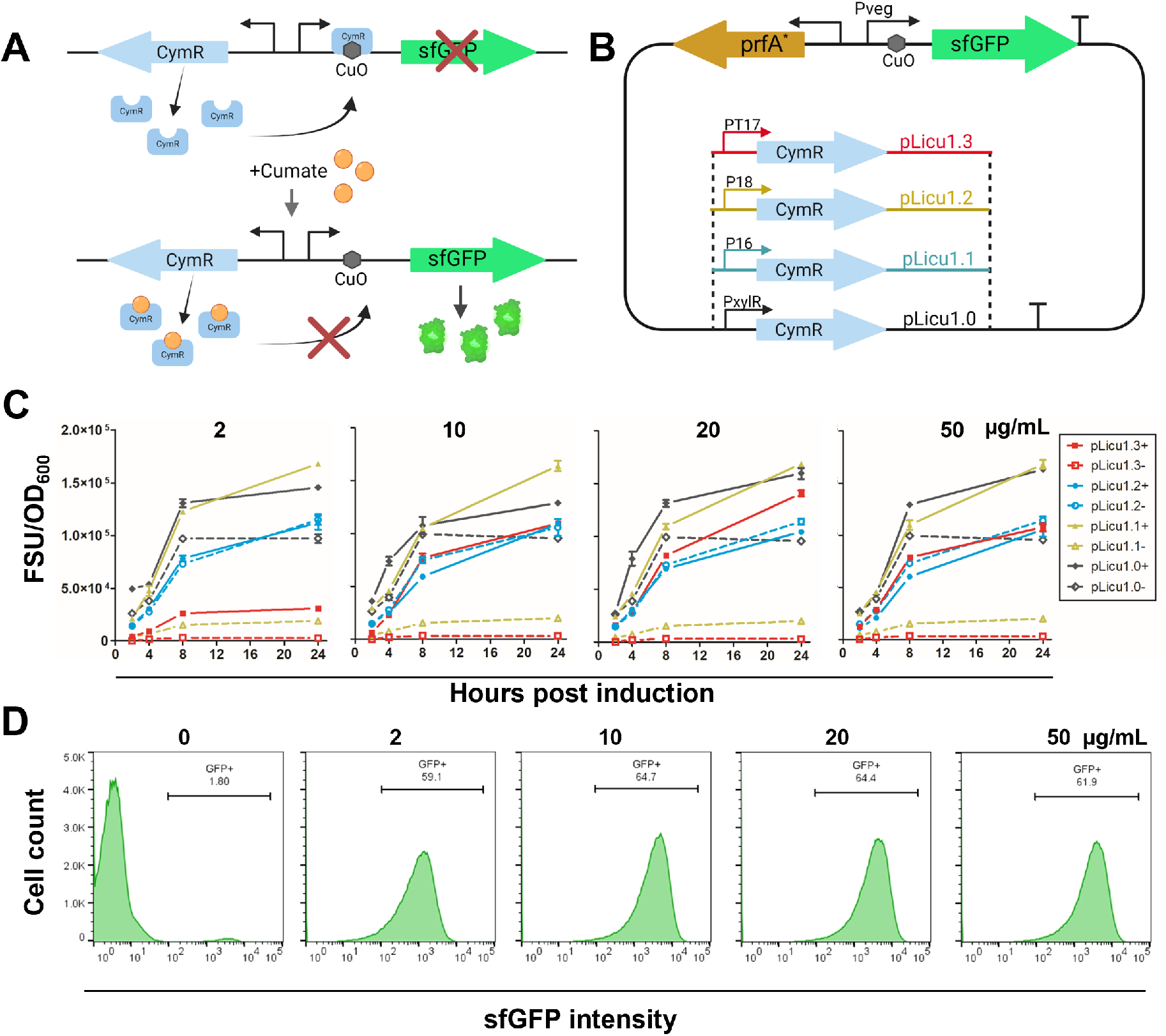
Engineering and testing of the cumate-inducible system in *Lm in vitro*. (**A**) Schematic of the cumate-inducible gene expression system in *Lm*. (**B**) Optimizing the *in vitro* cumate-inducible system. PxylR in pLicu1.0 was replaced with constitutive promoters (P16, P18, and PT17) to create pLicu1.1, 1.2, and 1.3. (**C**) The expression of sfGFP *in vitro. Lm* XFL-7.1 containing the *in vitro* cumate-inducible constructs was induced with 0–50 µg/mL cumate. sfGFP was measured at 2, 4, 8, and 24 h post-induction. FSU, fluorescence standard unit. Data represent mean ± SD of two biological replicates. (**D**) Flow cytometry analysis of sfGFP expression in *Lm. Lm* XFL-7 harboring pLicu1.3 was grown with 0–50 µg/mL cumate for 24 h followed by washing with PBS and measurements using the Guava EasyCyte 12HT flow cytometer. Data were analyzed using FlowJo™ v10.8 Software. Schematics are created using Biorender.com.

While suitable for gene regulation in *Lm in vitro*, pLicu1.3 is not adequate for regulating gene expression in intracellular *Lm* due to the lack of trLLO. To overcome toxicity of the CymR plus trLLO combination in *E. coli*, we expressed the *cymR* gene in the *Lm* chromosome from the PT17 promoter. To this end, PT17-CymR was cloned into vector pIMK2 that was integrated in the XFL-7 chromosome. The CuO-sfGFP construct was expressed from a strong, P_veg_ promoter^24^, in the therapeutic plasmid pGG-34 containing *trLLO* (Fig. 2A, pLicu2.0). The chromosomal CymR expression from the integrated pIMK2 plasmid appeared to be sufficient for repression of the plasmid-encoded CuO-sfGFP in the absence of cumate (Fig. 2A, pLicu2.0L). However, pLicu2.0L had a weak ribosome binding site (RBS) (strength, 4K^27^) upstream of sfGFP. To increase sfGFP expression, we engineered medium-(23K) and high-strength (96K) RBSs (Fig. 2A, pLicu2.0M, and pLicu2.0H). Since these constructs did not result in leaky expression, pLicu2.0H was used in subsequent experiments (Fig. 2B). The sfGFP levels expressed from pLicu2.0H were induced by 80-fold at 20 µg/mL cumate, making it a suitable for testing *in vivo* (Fig. 2B). Notably, at 24 h post-induction, sfGFP fluorescence decreased likely due to cell lysis and the release of sfGFP into the growth media. Flow cytometry analysis confirmed that pLicu2.0H was highly responsive to the addition of cumate, with 94 % *Lm* in liquid cultures being induced after 8 h at low, 10 µg/mL cumate cooncentrration (Fig. 2C). Compared to pLicu1.3, pLicu2.0H exhibited a broader dynamic range at lower cumate concentrations and was more responsive to cumate at earlier time points (Fig. 2B, C).

**Fig. 2.**
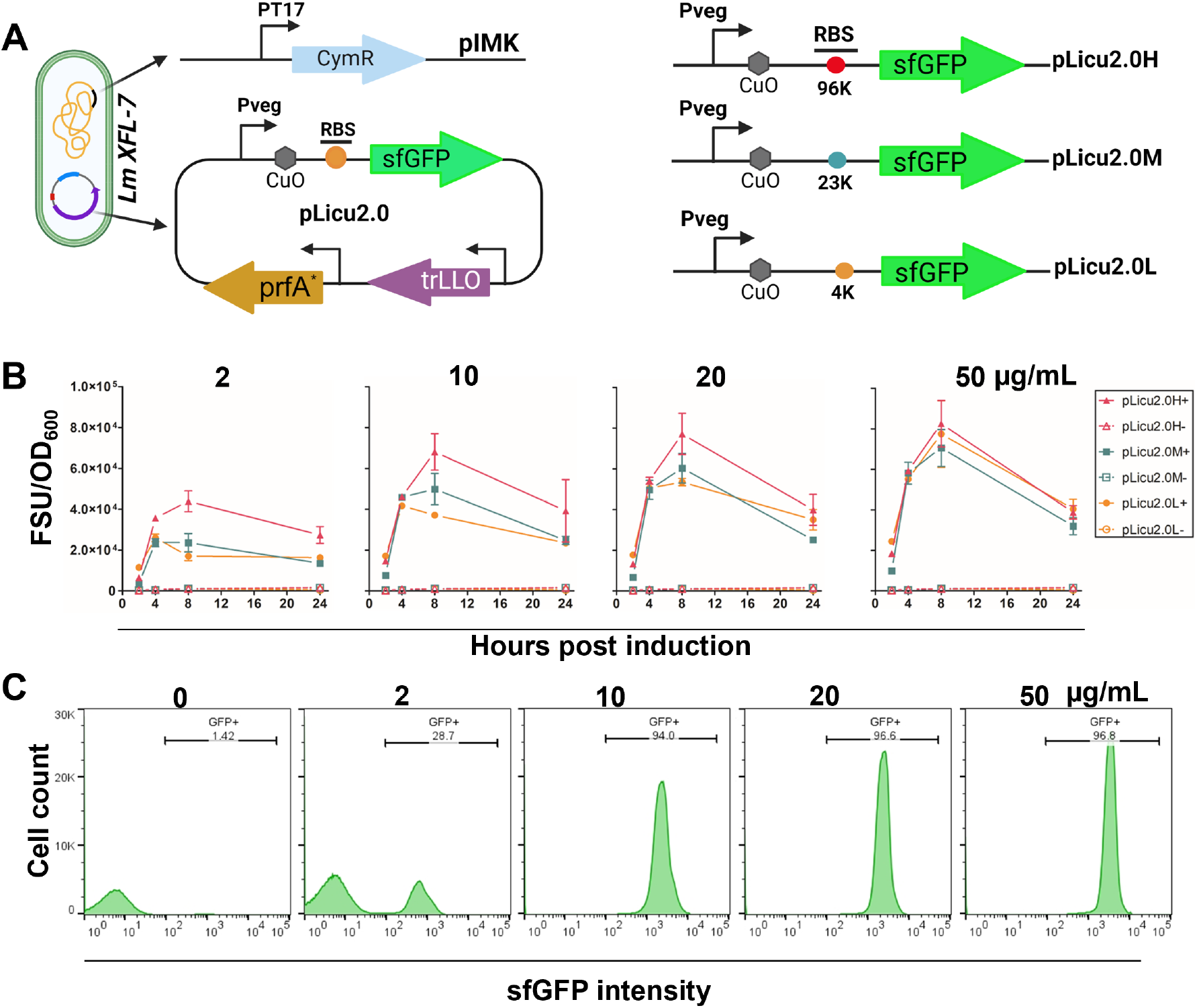
Engineering and testing of the cumate-inducible system for the use in *Lm in vivo*. (**A**) Schematic of the split *in vivo* cumate-inducible system. PT17-CymR was integrated into the chromosome via pIMK2, while the rest of the system is cloned in plasmid pGG-34. Three constructs with different RBS strengths (4K, 23K, and 96K) were tested, i.e., pLicu2.0L, 2.0M, and 2.0H. (**B**) sfGFP fluorescence in *Lm* XFL-7 grown at different cumate concentrations (0– 50 µg/mL) after 2, 4, 8, and 24 h post-induction. Data represent mean ± SD from two biological replicates. (**C**) Flow cytometry analysis of sfGFP expression in *Lm* XFL-7 harboring pLicu2.0H. *Lm* was grown with 0–50 µg/mL cumate for 8 h followed by washing with PBS and measuring using Guava EasyCyte 12HT flow cytometer. Data were analyzed using FlowJo™ v10.8 Software. Schematics are created using Biorender.com.

Next, we tested if cumate can induce gene expression in intracellular *Lm* inside the tumor cells (Fig. 3A). The 4T1 mouse breast cancer cells were infected with *Lm* strain carrying the chromosomal *cymR* gene and the pGG-34-derived plasmid containing CuO-sfGFP. The expression of sfGFP in intracellular *Lm* in cell culture was dependent on cumate concentration and RBS upstream of sfGFP, similar to what was observed in *Lm* cultures (Fig. 3A). The expression of sfGFP increased from 0 to 24 h post-induction and plateaued after 24 h (Fig. 3A). Consistent with our expectations, induction in intracellular *Lm* was slower and required higher cumate concentrations, in comparison to *Lm* cultures. Microscopy analysis confirmed sfGFP induction in intracellular *Lm* (Fig. 3B). To visualize intracellular *Lm*, a red-fluorescent protein, mScarlet-I, was cloned into pIMK-CymR under a constitutive promoter, pIMK-CymR-red, and this plasmid was integrated in the *Lm* chromosome. The microscopy images showed that cumate induced sfGFP expression in intracellular *Lm*, with higher cumate concentrations leading to higher expression (Fig. 3B), which is consistent with bulk sfGFP fluorescence data (Fig. 3A).

**Fig. 3.**
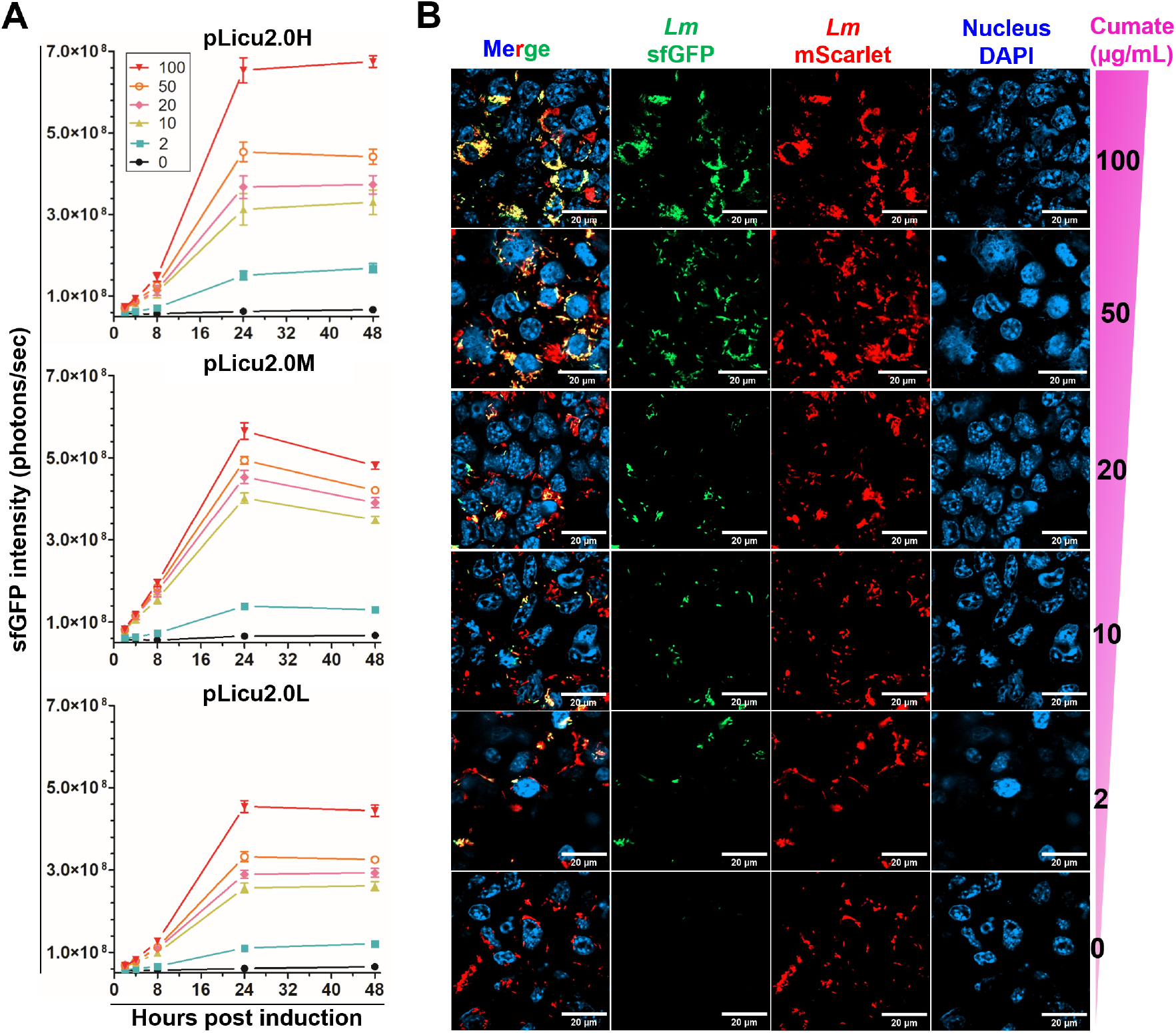
The cumate-inducible system tightly controls sfGFP expression in intracellular *Lm* in cell culture. (**A**) Gene induction in intracellular *Lm* in 4T1 breast cancer culture. 4T1 cells were infected with *Lm* XFL-7::pIMK-CymR harboring pLicu2.0 constructs at MOI 10 for 15 h prior to the addition of cumate (0–100 µg/mL). The expression of sfGFP in intracellular *Lm* was measured at 2, 4, 8, 24, and 48 h using IVIS with a plate mode; 10-sec exposure, 10% excitation power. Data represent mean ± SD from eight biological replicates. (**B**) Microscopy imaging of cumate-induced sfGFP expression in intracellular *Lm*. 4T1 cells were infected with *Lm* XFL-7::pIMK-CymR-red harboring pLicu2.0H. The cultures were induced with cumate (0–100 µg/mL) for 24 h, fixed with 4 % PFA, and imaged via confocal microscopy. Red, *Lm* expressing mScarlet; green, *Lm* expressing sfGFP; yellow, *Lm* expressing both mScarlet and sfGFP; blue, 4T1 cell nuclei.

*Lm* XFL-7 harboring plasmid pGG-34 propagates in TMEs to high levels after being eliminated from the healthy, non-tumor, tissues. To determine if the introduced genetic modifications affected strain behavior *in vivo*, the modified strain, designated XFL-7.1 (XFL-7::pIMK-CymR-red carrying pLicu2.0H), was injected in 4T1-breast tumors of female BALB/c mice. XFL-7.1 colonized tumor-bearing mice similarly to the original XFL-7 strain^4,11^. XFL-7.1 numbers in tumors reached levels that were 2-to 4 orders of magnitude higher than in other organs on days 1 and 3 post-infection (Fig. 4A). As expected, XFL-7.1 was rapidly eliminated from the healthy organs, but persisted inside tumors for longer period of time (Fig. 4A). These findings suggest that introduction of the cumate-inducible system did not affect *Lm* behavvior *in vivo*.

**Fig. 4.**
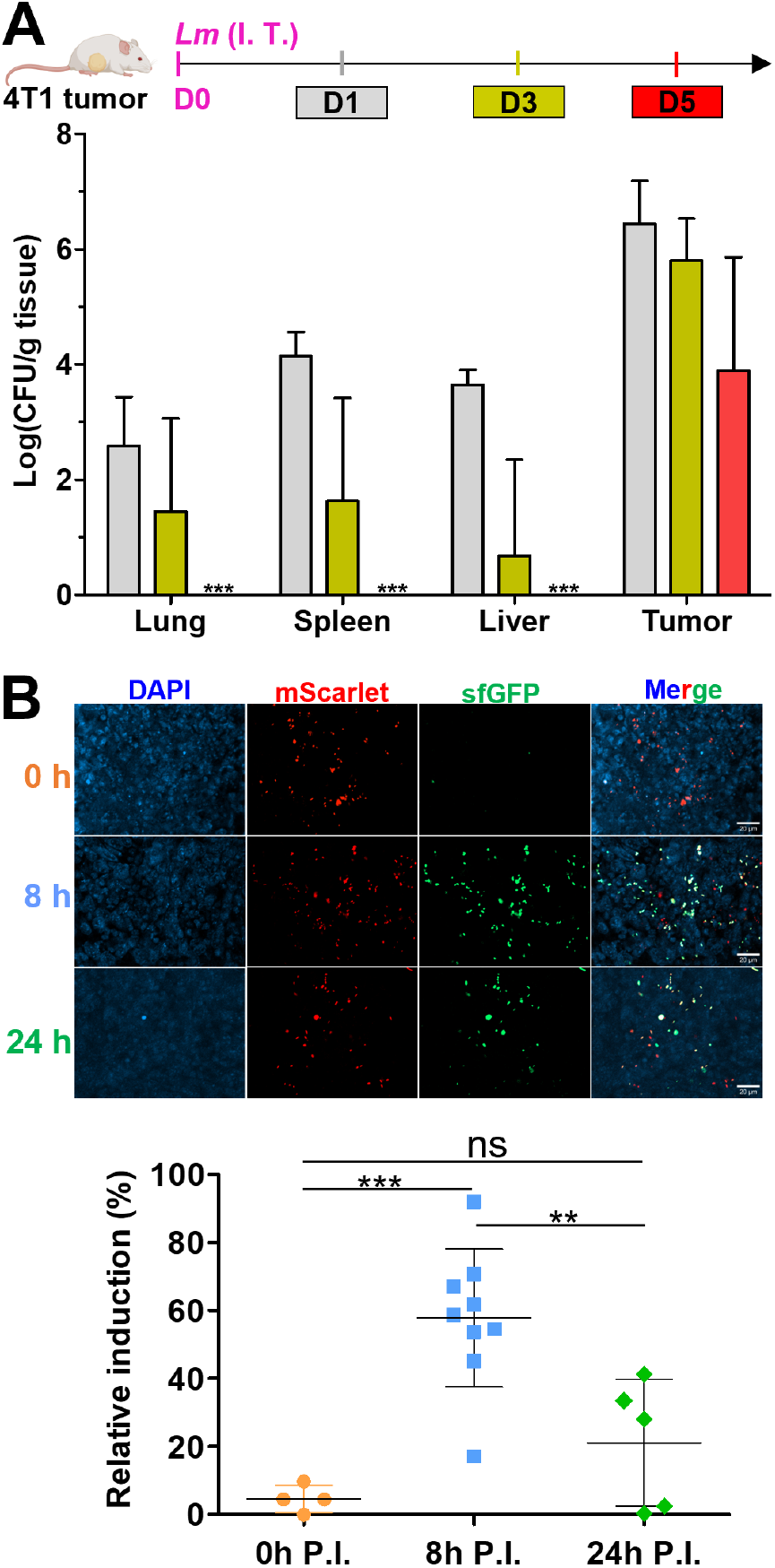
The cumate-inducible system robustly induces sfGFP expression in intracellular *Lm* intratumorally. 4T1 breast tumor cells were implanted into the mammary fat pads of female BABL/c mice. *Lm* XFL-7.1 was injected into the tumors when they reached 50–100 mm^3^. (**A**) The accumulation of *Lm* XFL-7.1 in tissues of the 4T1 tumor-bearing mice. The lung, liver, spleen, and tumor tissues were collected and homogenized on days 1, 3, and 5 post *Lm* XFL-7.1 injection. *Lm* CFUs were counted by plating of the diluted tissue homogenates on BHI agar supplemented with 10 µg/mL chloramphenicol. ***, below detection. Data represent mean ± SD from six mice per group. (**B**) The induction of sfGFP expression in intratumoral *Lm* XFL-7.1. On day 3 post *Lm* injection, intratumoral *Lm* XFL-7.1 was induced with 3 mg/mouse of water-soluble cumate delivered via intraperitoneal injection. Tumors were isolated at 0, 8, and 24 h post-induction, fixed with 4% PFA followed by sectioning. Relative induction percentage was calculated based on Manders’ coefficient using ImageJ with the JACoP plugin. One dot represents one data point from a field of view of 6 tumors per group. Statistical significance was calculated by one-way ANOVA with a Tukey’s multiple comparison tests using Prism 5. P < 0.05, ns: not significant. Data represent mean ± SD from six mice per group.

To visualize gene induction in intracellular *Lm* in tumors, 4T1 tumors were injected with *Lm* XFL-7.1. On day 3 post-injection, when intracellular *Lm* numbers in TMEs peak, mice were intraperitoneally administered 3 mg/mouse of water-soluble cumate. This dose was based on the report that 1.5 mg/mouse of water-soluble cumate induced expression of virus-delivered genes in mouse livers^31^. Our attempts to image sfGFP in the tumors of live animals failed likely because *Lm* numbers were insufficient (∼10^6^ CFU/g tumor). The extracellular bacteria, *E. coli* and *Salmonella*, typically reach 10^9^–10^10^ CFU/g tumor, which is sufficient for whole-animal GFP imaging^6,8^. Therefore, to visualize sfGFP induction, mice were euthanized at 0, 8, 24 h post-induction, and tumors were isolated, fixed, and sectioned for microscopy. Robust induction of intracellular *Lm* inside tumors was observed at 8 h post induction with approximately 60 % of *Lm* bacteria expressing sfGFP, which represents at least a 10-fold increase compared to 0 h (Fig. 4B). The percentage of induced *Lm* significantly decreased at 24 h post-induction, suggesting that sfGFP overexpression lowers *Lm* vitality inside tumor cell. This observation suggests that induction of antitumor payloads *in vivo* need to be balanced with intracellular *Lm* survival. Further experiments will be required to evaluate how orally delivered cumate affects gene induction in intratumoral *Lm*.

In conclusion, the developed here cumate-dependent system enables tightly regulated gene induction in *Lm in vitro*, in *Lm*-infected cell cultures and in the intracellular *Lm* in infected tumors of live animals. This system is anticipated to enhance the efficacy and safety of *Lm*-based cancer therapies.

## Abbreviations

CFU: colony-forming unit
IVIS: *in vivo* imaging system
*Lm*: *Listeria monocytogenes*
MOI: multiplicity of induction
*prfA*: gene encoding the *Lm* master virulence regulator (positive regulatory factor A)
RBS: ribosome-binding site
TAA: tumor-associated antigen
TME: tumor microenvironment
trLLO: truncated listeriolysin O

## Acknowledgments

We thank Dr. Claudia Gravekamp for providing strain XFL-7 and plasmid pGG34, Drs. Zhaojie Zhang and Qian-Quan Sun for assistance with microscopy and tissue sectioning. This work was supported by National Cancer Institute (NCI) Grant R21 CA249814. The Integrated Microscopy Core Facility is supported by NIGMS P20 GM121310.

