## Supporting Information for "Robust inducible gene expression in intracellular *Listeria monocytogenes in vivo*"

Contents:

Suppl. Fig. 1. Instability of the plasmids carrying the *cymR* and *trLLO* genes.

Suppl. Fig. 2. Assessment of cumate toxicity to *Lm*.

Suppl. Fig. 1. **Instability of the plasmids carrying the *cymR* and *trLLO* genes.** (A) Electrophoresis analysis showing instability of the pGG-34 plasmids expected to contain the cumate-induction system. Plasmids were isolated from multiple *E. coli* NEB Stable colonies. (B, C) Plasmid pLicu1.0 lacking *trLLO* is stable in *E. coli* (B) and in *Lm* XFL-7 (C).

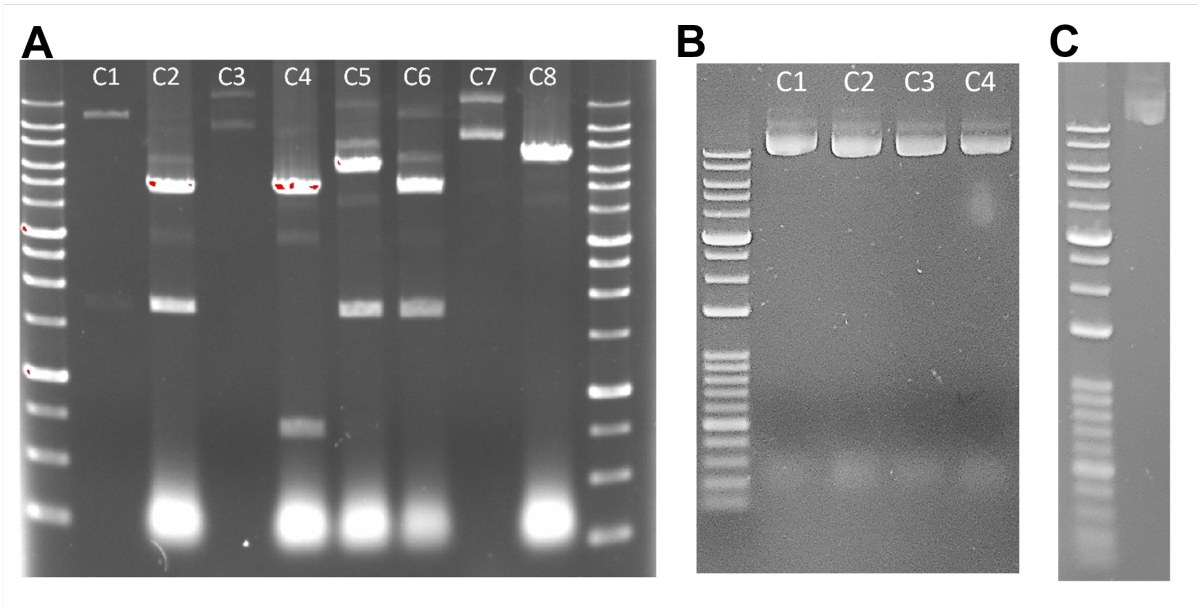


Suppl. Fig. 2. **Assessment of cumate toxicity to *Lm*.** The *Lm* cultures were grown in BHI broth supplemented with at 0–500 µg/mL cumate for 24 h. The cultures were diluted and plated on BHI agar to count CFUs. Shown are means ± SD of three technical replicates.

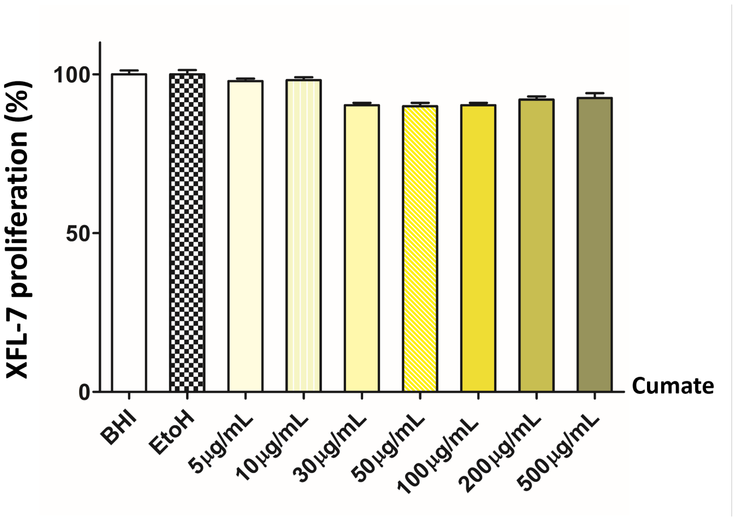
